# GSK3 inhibition reverts mesenchymal transition in human primary corneal endothelial cells

**DOI:** 10.1101/2022.11.25.517972

**Authors:** Eleonora Maurizi, Alessia Merra, Claudio Macaluso, Davide Schiroli, Graziella Pellegrini

## Abstract

Human corneal endothelial cells are organized in a tight mosaic of hexagonal cells and serve a critical function in maintaining corneal hydration and clear vision. Regeneration of the corneal endothelial tissue is hampered by its poor proliferative capacity, which is partially retrieved *in vitro*, albeit only for a limited number of passages before the cells undergo mesenchymal transition (EnMT). Although different culture conditions have been proposed in order to delay this process and prolong the number of cell passages, EnMT has still not been fully understood and successfully counteracted. In this perspective, we identified herein a single GSK3 inhibitor, CHIR99021, able to revert and avoid EnMT in primary human corneal endothelial cells (HCEnCs) from old donors until late passages *in vitro* (P8), as shown from cell morphology analysis (circularity). In accordance, CHIR99021 reduced expression of α-SMA, an EnMT marker, while restored endothelial markers such as ZO-1, Na^+^/K^+^ ATPase and N-cadherin, without increasing cell proliferation. A further analysis on RNA expression confirmed CHIR99021 induced downregulation of EnMT markers (αSMA and CD44), upregulation of the proliferation repressor p21 and revealed novel insights into the β-catenin and TGFβ pathways intersections in HCEnCs. The use of CHIR99021 sheds light on the mechanisms involved in EnMT and brings a substantial advantage in maintaining primary HCEnCs in culture until late passages, while preserving the correct morphology and phenotype. Altogether, these results bring crucial advancements towards the improvement of the corneal endothelial cells based therapy.

## Introduction

The innermost layer of the cornea, the corneal endothelium (CE), efficiently transports solutes from and to the aqueous humor, thus maintaining corneal transparency and function. Corneal endothelial cells (CEnCs) are poorly mitotic *in vivo*^*1*^, as a consequence of the sealed tight cell-cell junctions and the presence of proliferative inhibitors in the aqueous humor^2-3^. Cell migration and enlargement^1, 4^ are the mechanisms that counteract a constant decrease in cell density (0.6% every year)^5-6^; nevertheless exogenous factors, such as surgical interventions, infections or physical trauma, and corneal diseases may irreversibly damage the CE, thus impairing this delicate equilibrium. In this case, an altered CE becomes unable to play its function^4^, leading to corneal opacification and, in the last instance, to blindness. To date, transplantation is the only accepted treatment to cure irreversibly damaged corneas, for which CE dysfunction represents the most frequent indication^7^. Although quite diffused and generally safe, corneal grafts are not equally accessible worldwide and still present a relevant failure rate^7^.

Since CEnCs are able to acquire, albeit in a limited manner, a proliferative capacity in culture, researchers are willing to promote CE regeneration directly *in vivo* or *in vitro*^*7*^, prior to transplantation.

In the first-in-human clinical trial, Kinoshita’s group successfully transplanted human (H)CEnCs obtained from young donors and expanded for a maximum of three passages in culture^8^. Nevertheless, several improvements are still necessary since HCEnCs easily alter their morphology and function *in vitro*^9^. This impairment, which is exacerbated in cells from older donors, drastically increases with passages and is mainly a consequence of mesenchymal transformation (endothelial to mesenchymal transition, EnMT)^10^, a cellular process occurring also in cancer cells, and during normal expansion of several stem cells^11^. In line with this observation, EnMT might represent a physiological and reversible process that naturally underlies HCEnCs migration and proliferation. However, if EnMT is dysregulated, for instance during *in vitro* expansion when HCEnCs are induced to leave their quiescence (G0/G1) and become proliferative (G2), it may lead to an irreversible mesenchymal transformation^10^. EnMT transformed HCEnCs (positive for specific markers such as α-SMA), arise not only after passaging^12^, but also consequently to the cell junctions disruption following EDTA treatment^13^, as well as upon growth factors stimulation^13-17^. It has been largely documented that epithelial and fibroblast growth factors (EGF and FGF, respectively) are able to promote cellular proliferation while preserving the cellular phenotype^18-20^, but only for few passages^13-17, 21^, while transforming growth factor (TGFβ) induces EnMT upon certain conditions^17^. HCEnCs undergoing a mesenchymal transformation present with an altered cellular functionality, mainly caused by the loss of cellular structure (in terms of polarity, motility, extracellular matrix production, cytoskeleton modifications and loss of cellular contacts)^10^. As a consequence, EnMT transformed HCEnCs are not suitable for developing a cell therapy as they are not able to reproduce an intact and functional endothelium, but they can induce tissue fibrosis^21^. Although the maximum number of passages that HCEnCs reach in culture depends primarily on donor characteristics, EnMT represents one of the main hurdles limiting *in vitro* expansion. Aiming at increasing the maximum number of passages, researchers have investigated some alternative culture media conditions to promote proliferation and stem at the same time EnMT undesired effects.^7, 22-24^. In particular TGFβ^24^ and ROCK pathway inhibition^25^ and, more recently, a dual media approach (the first during the proliferative state and the second of maintenance)^22, 26-28^ have found a large consensus^7, 9^ as valid solutions to preserve HCEnCs phenotype. Nevertheless, no key mechanisms able to revert the EnMT and possible approaches to reach this aim have been successfully proposed. Some hints support the hypothesis that the β-catenin pathway might be a reasonable target: i) The Wnt/β-catenin pathway is finely regulated in other cell types to switch from a proliferative to a differentiate state^29-30^. ii) Dysregulation of this pathway is an important player in mesenchymal transformation^31-32^. iii) β-catenin involvement was previously linked to this delicate balance from us and other groups in HCEnCs^13, 16, 33-34^. Moreover, we recently proved that β-catenin pathway is finely tuned when CEnCs are dissected and expanded *in vitro*^16^. This pathway resulted fundamental to maintain proliferation, while its activation through a small molecule CHIR99021 was not able to increase cellular proliferation. We also measured the α-SMA positive cells at low passage in rabbit primary CEnCs and we found a small but not significant decrease of this parameter. CHIR99021 has been largely used to induce cellular differentiation^35-37^, acting on the β-catenin stability. In corneal endothelium it promotes the direct transformation of corneal stroma precursors cells to corneal endothelial cells^38^ and recently Wang and collaborators showed that it was able to counteract the TGFβ1 induced cell modification, in particular α-SMA overexpression, in a cell line of corneal endothelium^39^. Starting from these pieces of evidence, we demonstrate here the CHIR99021 capability to both avoid and revert EnMT in primary HCEnC from old donor corneas during *in vitro* expansion, up to high passages (P8).

## Methods

### Ethical statement

Donor human corneas from Italian Eye Banks, unsuitable for transplantation, were obtained with their relatives written consent for research use. The tissues were handled in accordance with the declaration of Helsinki. The experimental protocol was approved by ISS-CNT (Italian National Transplant Centre) and by the local ethical committee (Comitato Etico dell’Area Vasta Emilia Nord, p. 0002956/20).

### Primary HCEnCs culture

Human corneal tissues were preserved in Eusol at 4°C and used within 15 days from the donor’s death; a list of details for each cornea used and relative experiments is shown in Table1. Except from one donor who was 26 and another who was 51 years old, the majority ranged from 63 to 81 years old. HCEnCs were isolated through Descemet’s stripping and digestion with 1.5 mg/ml Collagenase A (Roche) for 3 hours (h) at 37 °C. After 5 minutes (min) in TrypLE (Thermo Fisher Scientific) at 37°C, HCEnCs were pelleted at 1,200 rpm for 3 min and plated on FNC Coating mix (AthenaES) treated wells.

**Table 1.**
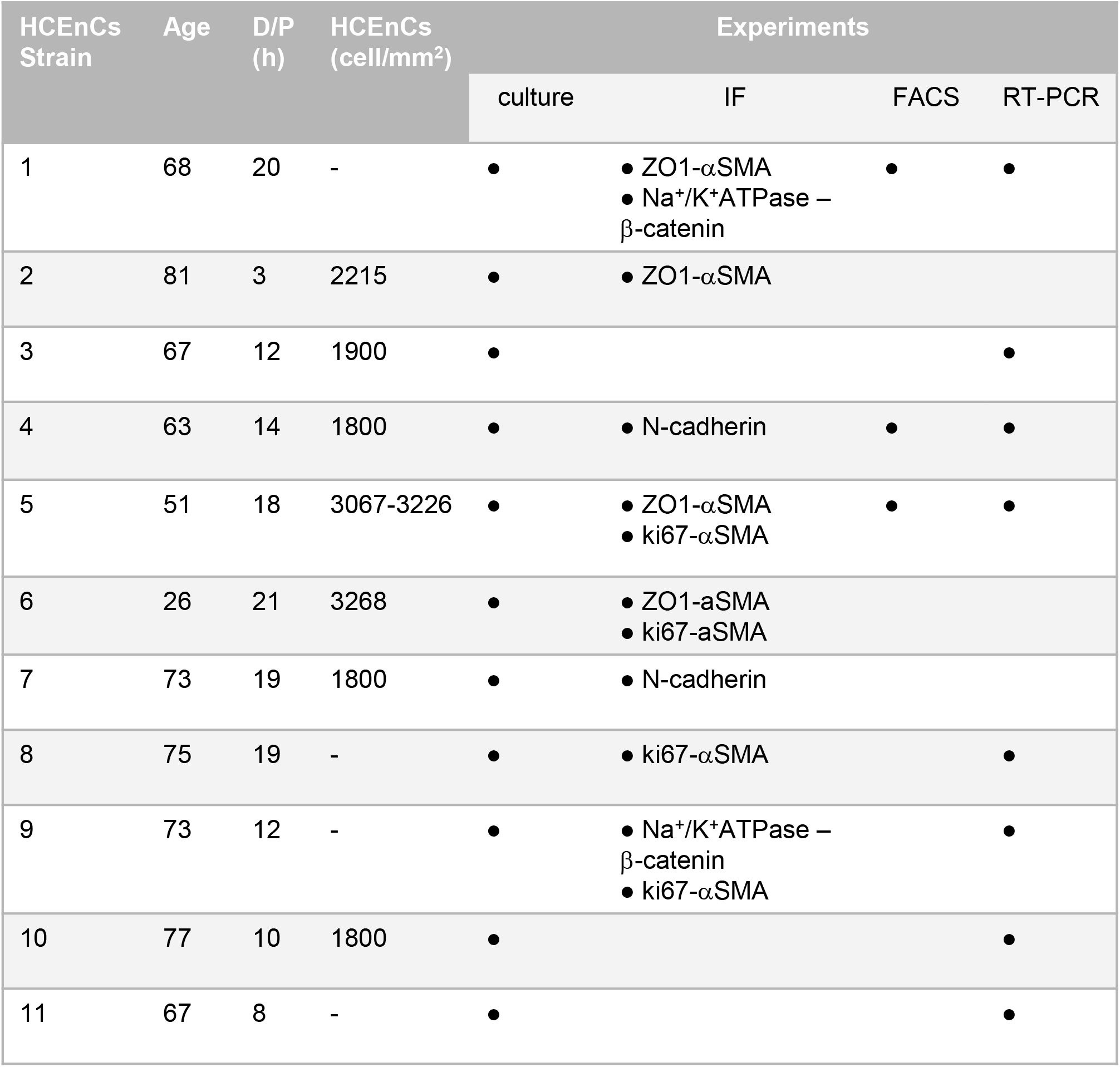
Donor human corneas used for the experiments. D/P indicates the interval (in hours) between the death and the preservation of donor human corneas. HCEnCs (cell/mm^2^) indicates the HCEnCs count done by the eye bank at the time of preservation. On the right, the columns indicate the experiments done for each strain: culture, immunofluorescence analysis (IF), FACS analysis, qRT-PCR.

| HCEncs Strain | Age | D/P (h) | HCEncs (cell/mm<sup>2</sup>) | Experiments |  |  |  |
| --- | --- | --- | --- | --- | --- | --- | --- |
|  |  |  |  | culture | IF | FACS | RT-PCR |
| 1 | 68 | 20 | - | ● | ● ZO1-αSMA<br>● Na<sup>+</sup>/K<sup>+</sup>ATPase – β-catenin | ● | ● |
| 2 | 81 | 3 | 2215 | ● | ● ZO1-αSMA |  |  |
| 3 | 67 | 12 | 1900 | ● |  |  | ● |
| 4 | 63 | 14 | 1800 | ● | ● N-cadherin | ● | ● |
| 5 | 51 | 18 | 3067-3226 | ● | ● ZO1-αSMA<br>● ki67-αSMA | ● | ● |
| 6 | 26 | 21 | 3268 | ● | ● ZO1-aSMA<br>● ki67-aSMA |  |  |
| 7 | 73 | 19 | 1800 | ● | ● N-cadherin |  |  |
| 8 | 75 | 19 | - | ● | ● ki67-αSMA |  | ● |
| 9 | 73 | 12 | - | ● | ● Na<sup>+</sup>/K<sup>+</sup>ATPase – β-catenin<br>● ki67-αSMA |  | ● |
| 10 | 77 | 10 | 1800 | ● |  |  | ● |
| 11 | 67 | 8 | - | ● |  |  | ● |

HCEnCs were cultured at 37°C in 5% CO_2_, changing the growth medium every other day. The growth medium is composed of OptiMEM-I (Thermo Fisher Scientific), 8% HyClone fetal bovine serum (FBS; Fisher Scientific), 5 ng/mL epidermal growth factor (EGF; Thermo Fisher Scientific), 20 μg/mL ascorbic acid (Sigma-Aldrich), 200 mg/L calcium chloride (Sigma-Aldrich), 0.08% chondroitin sulphate (C4384, Sigma-Aldrich), and penicillin/streptomycin (Euroclone). Upon confluency, HCEnCs were rinsed in DPBS and passaged at a ratio of 1:2 or 1:3 with TrypLE (Thermo Fisher Scientific) for 10-15 min at 37°C in 5% CO_2_. Sub-confluent cultures were harvested 24h after plating.

CHIR99021 (SML1046, Sigma-Aldrich, concentration range of 0.1-10 μM) and TGFβ inhibitor (TGFβI, SB431542, Sigma-Aldrich, 1 μM) were added to the culture media and replaced at any medium change.

### Immunofluorescence

Immunofluorescence staining was performed on primary cultured HCEnCs after fixation in 3% PFA, 15 min at room temperature (RT). Triton X-100 (Bio-Rad) used at 1% for 10 min at RT allowed cell permeabilization and a solution of bovine serum albumin (BSA; Sigma-Aldrich) 2%, FBS 2%, Triton X-100 at 0.01% in PBS was used for 30 min at 37°C to block the non-specific binding sites. Primary and secondary antibodies were incubated for 1 h at 37°C while nuclei were counterstained with DAPI (Roche) at RT for 5 min before mounting the glass coverslips using DAKO mounting medium (Agilent). Primary antibodies used herein are listed in Table 2 while secondary antibodies used are: Alexa Fluor 488 anti-rabbit, 1:2000, and Alexa Fluor 568 anti-mouse, 1:1000 (Thermo Fisher Scientific). Images were obtained with a confocal microscope (LSM900 Airyscan—Carl Zeiss).

**Table 2.** Antibodies and relative dilutions used for IF analysis.

| protein | reference | dilution |
| --- | --- | --- |
| ZO-1 | 40-2200 (Thermo Fisher) | 1:100 |
| α-SMA | A5228 (Sigma Aldrich) | 1:200 |
| β-catenin | ab32572 (Abcam) | 1:100 |
| Na<sup>+</sup>/K<sup>+</sup> ATPase | ab7661 (Abcam) | 1:200 |
| N-cadherin | 610920 (BD) | 1:100 |
| ki67 | ab15580 (Abcam) | 1:50 |

Quantification of α-SMA staining was obtained by counting the number of positive cells (primary antibody signal), relative to the total number of cells in that field (DAPI staining). The values obtained from different strains were expressed in percentage with standard deviation (three fields for each replicate were collected).

CellEvent® Caspase 3/7 Green (Thermo Fisher, UK) was used to evaluate cell apoptosis, following the manufacturer’s instructions. As a positive control for this assay, HCEnCs were treated with 10 mM H_2_O_2_ for 2 h.

### Cell Circularity

Cell circularity was determined using ImageJ software: morphometric values of the perimeter and area of cells were obtained from phase contrast images of the culture by manually outlining each cell borders, as shown in previously published papers^22^. An average of 50 cells for each condition were analysed, from three different cell strains each (n=3).

A perfect circle has a circularity value of 1: polygonal HCEnCs have a value closer to 1 if compared with HCEnCs with an elongated fibroblastic morphology, which circularity is closer to zero.

### Cell cycle analysis by FACS

The cell cycle was studied by staining sub-confluent cultures of HCEnCs with Propidium Iodide (PI; Sigma-Aldrich). HCEnCs suspension from cell culture was washed with DPBS and incubated for 1 h at 4°C in the dark with 250 μL of a PBS solution composed by PI 50 μg/mL, Triton X-100 (Bio-Rad, USA) at 0.1%. Cells were then analysed using BD FACS Canto II (BD BIOSCIENCES; San Jose, CA USA). For each sample, 20,000 events were considered for the analysis to ensure statistical relevance and results were analysed with a ModFit 3.0 software.

### RT-PCR

HCEnCs RNA was extracted by RNeasy plus Micro Kit (Qiagen), quantified with Nanodrop 100 (Thermo Fisher Scientific) and reverse transcribed by the High Capacity cDNA Reverse Transcription Kit (Thermo Fisher Scientific).

7900HT Fast Real-Time PCR System (Thermo Fisher Scientific) was used for RT-PCR assays, using the TaqMan Real Time PCR Assays probes and primers for SyBr Green listed in Table 3.

**Table 3.** Probes used for the qRT-PCR experiments. TaqMan assays were all purchased from Thermo Fisher Scientific.

| Gene | Probes - primers |
| --- | --- |
| p27 | Hs00153277_m1 |
| p21 | Hs00355782_m1 |
| PITX2 | Hs04234069_mH |
| SOX2 | Hs01053049_s1 |
| GAPDH | Hs02786624_g1 |

**Table 4.**
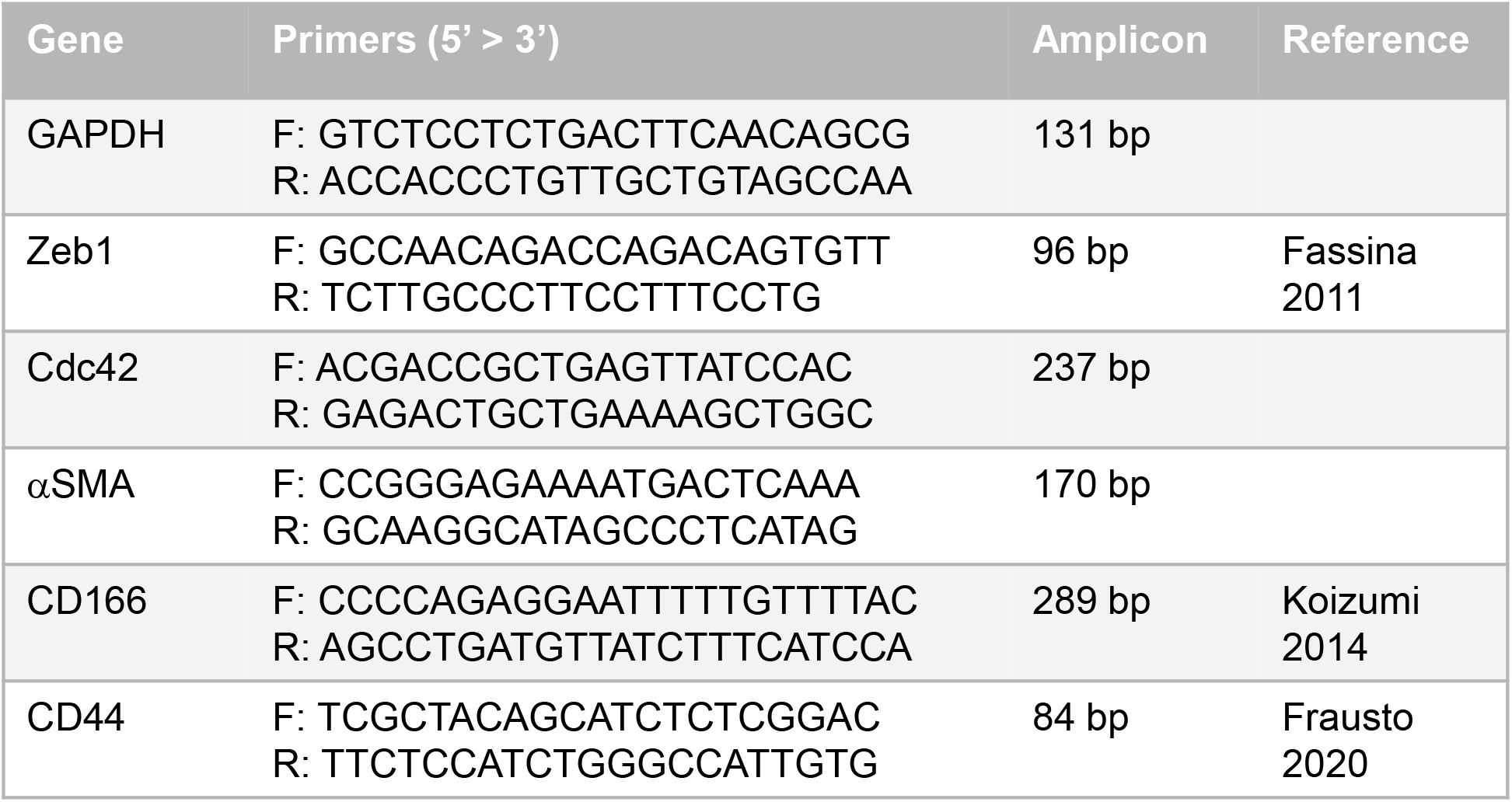
Primers used for the qRT-PCR experiments.

GAPDH was used as housekeeping control and ΔCt and ΔΔCt calculations were performed to evaluate effective RNA expression. Each gene was evaluated in three different strains for each condition, in two strains for the TGFβI analysis.

### Statistical analysis

Microsoft Excel 2010 and GraphPad Prism 5 software were used for data and statistical analysis. Values were represented as mean ± standard deviation (SD). Statistical comparison was done using two-tailed Student’s t-test, while gene expression data were compared with a Ratio Paired t-test. Significance was set at p < 0.05 and the number of replicates are indicated in each experiment.

## Results

### CHIR99021 (CHIR) regulates EnMT in primary cultures of HCEnCs

Primary HCEnCs were treated with CHIR99021 at different concentrations (1-3-10 μM) to evaluate its dose related effect upon EnMT (Figure 1).

**Figure 1.**
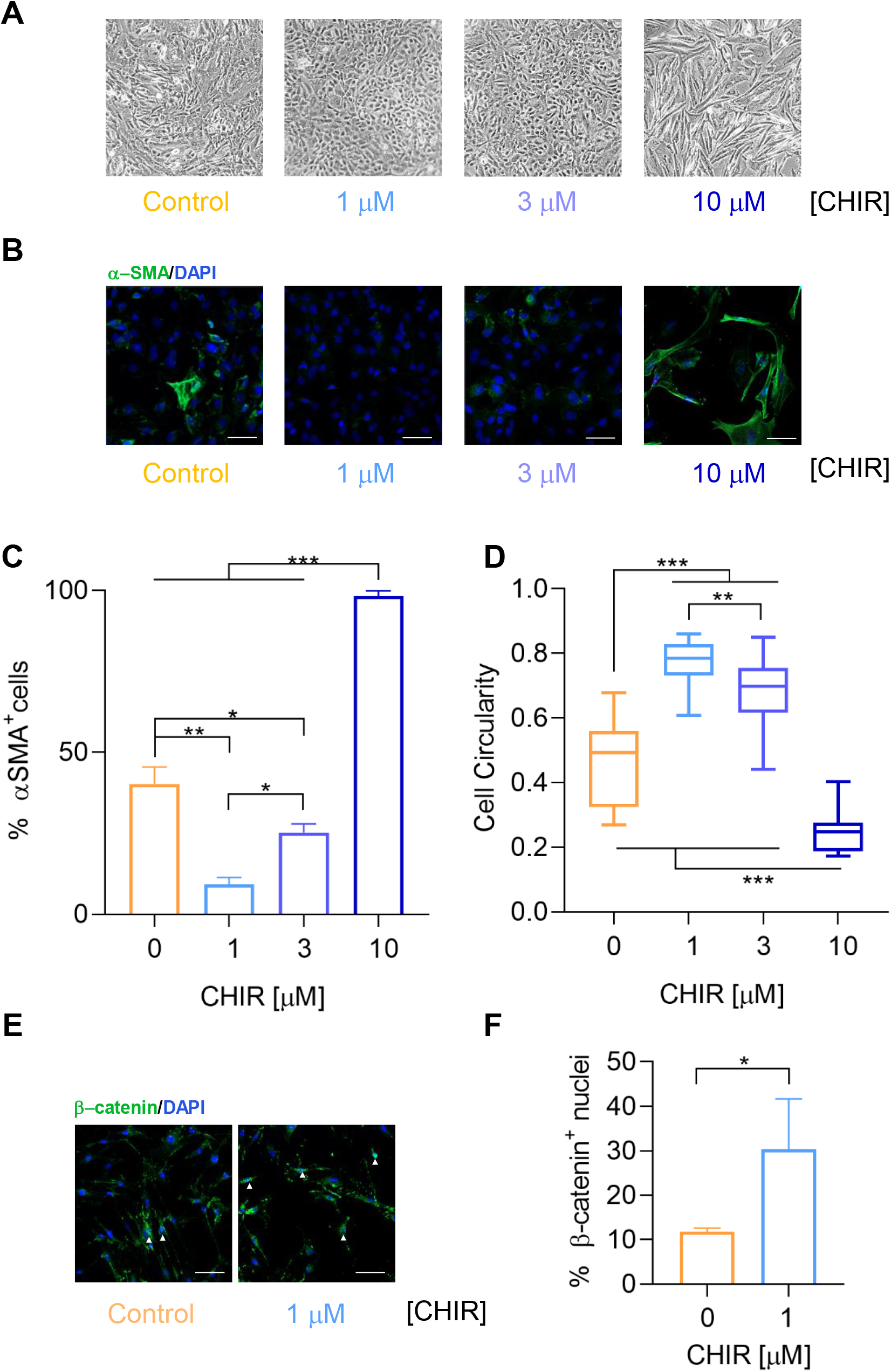
CHIR99021 titration in primary cultures of human corneal endothelial cells (HCEnCs). A) Representative phase-contrast images of primary HCEnCs culture at passage 1 (P1), showing the morphology differences between cells treated with CHIR99021 at different concentrations (1 - 3 - 10 μM) and control. B) Representative immunofluorescence microscopy images of HCEnCs at P1, showing expression of the α-SMA (green) EnMT marker on CHIR99021 treated and untreated HCEnCs. DAPI (blue) counterstains nuclei. Scale bar 50 μm. C) Quantification of the percentage of HCEnCs expressing α-SMA protein in samples treated with CHIR99021 at different concentrations (1 - 3 - 10 μM) were compared with their relative untreated control, as seen in Figure 1B. D) Box plot comparing cell circularity values between HCEnCs treated with CHIR99021 at different concentrations (1 - 3 - 10 μM) and their relative untreated control (n=3). E) Representative immunofluorescence microscopy images of HCEnCs at P4 showing expression of the β-catenin (green) on HCEnCs treated and untreated with CHIR99021 at 1 μM (CHIR). DAPI (blue) counterstains nuclei, white arrows indicate where β-catenin shows a nuclear signal. Scale bar 50 μm. F) Quantification of the percentage of HCEnCs expressing nuclear β-catenin protein in samples treated with CHIR, as compared with their relative untreated control, as seen in Figure 1E. Quantitative data are expressed as values ± SD. Statistical significance was assessed using a Student t-test and set at p<0.05. p values are indicated as following: ns (not significative) when p>0.05, * when p<0.05, ** when p<0.01, *** when p<0.001.

A clear variation of HCEnCs morphology was initially observed (Figure 1A): when CHIR99021 at 1-3 μM was added to the culture media, HCEnCs at passage 1 (P1) appeared as more polygonal if compared to the untreated control. An elongated morphology was observed instead whenever HCEnCs were treated with CHIR99021 at 10 μM. The immunocytochemistry analysis showed a reduced expression of α-SMA, marker of EnMT, in cells treated with CHIR99021 at 1-3 μM if compared to the control. CHIR used at 10 μM accelerated the process of EnMT, as observed from α-SMA expression that was increased in every cell (Figure 1B). Quantification of the percentage of HCEnCs expressing α-SMA in immunofluorescence images shown in Figure 1B demonstrates that the most effective CHIR99021 concentration, able to significantly reduce α-SMA expression to 9.2 ± 2.1% from 40.2 ± 5.2% of the control (p = 0.0046), was 1 μM (Figure 1C).

The maintenance of a hexagonal shape is a characteristic feature of HCEnCs and is fundamental for HCEnCs to exert their biological function^40-41^. For this reason, a morphological study on cell morphology from Figure 1A allowed evaluating the variation of HCEnCs circularity between the conditions under investigation (Figure 1D). HCEnCs treated with 1 μM CHIR99021 showed an average circularity value of 0.77, which is the closest to 1 (value of a perfect circle), if compared with the control (0.46), the 3 μM (0.69) and the 10 μM (0.25) CHIR99021 treated cells. CHIR99021 used at 1 μM was therefore chosen for all the subsequent experiments and will be indicated from now onwards as CHIR. As known from the literature, CHIR inhibits GSK-3β and thus stabilises intracellular β-catenin, as confirmed here in immunofluorescence analysis of HCEnCs at P4 (Figure 1E). β-catenin positive nuclei were quantified, resulting in an increase to 30.4 ± 6.5% in CHIR treated cells from a 11.8 ± 0.44% of the control (Figure1F).

CHIR99021 at 1 μM demonstrated here its capacity to reduce the process of EnMT in primary HCEnCs by restoring cell morphology and decreasing the expression of α-SMA marker. These results are crucial to counteract the EnMT process that hampers the opportunity for these cells to be amplified in culture and thus to be used for further studies or for clinical application.

### CHIR reverses the mesenchymal phenotype in primary cultures of HCEnCs

Primary HCEnCs, already presenting an elongated morphology (P3), were treated with CHIR for several subsequent passages in culture: CHIR demonstrated capable of reverting the mesenchymal phenotype at any passage (Figure 2).

**Figure 2.**
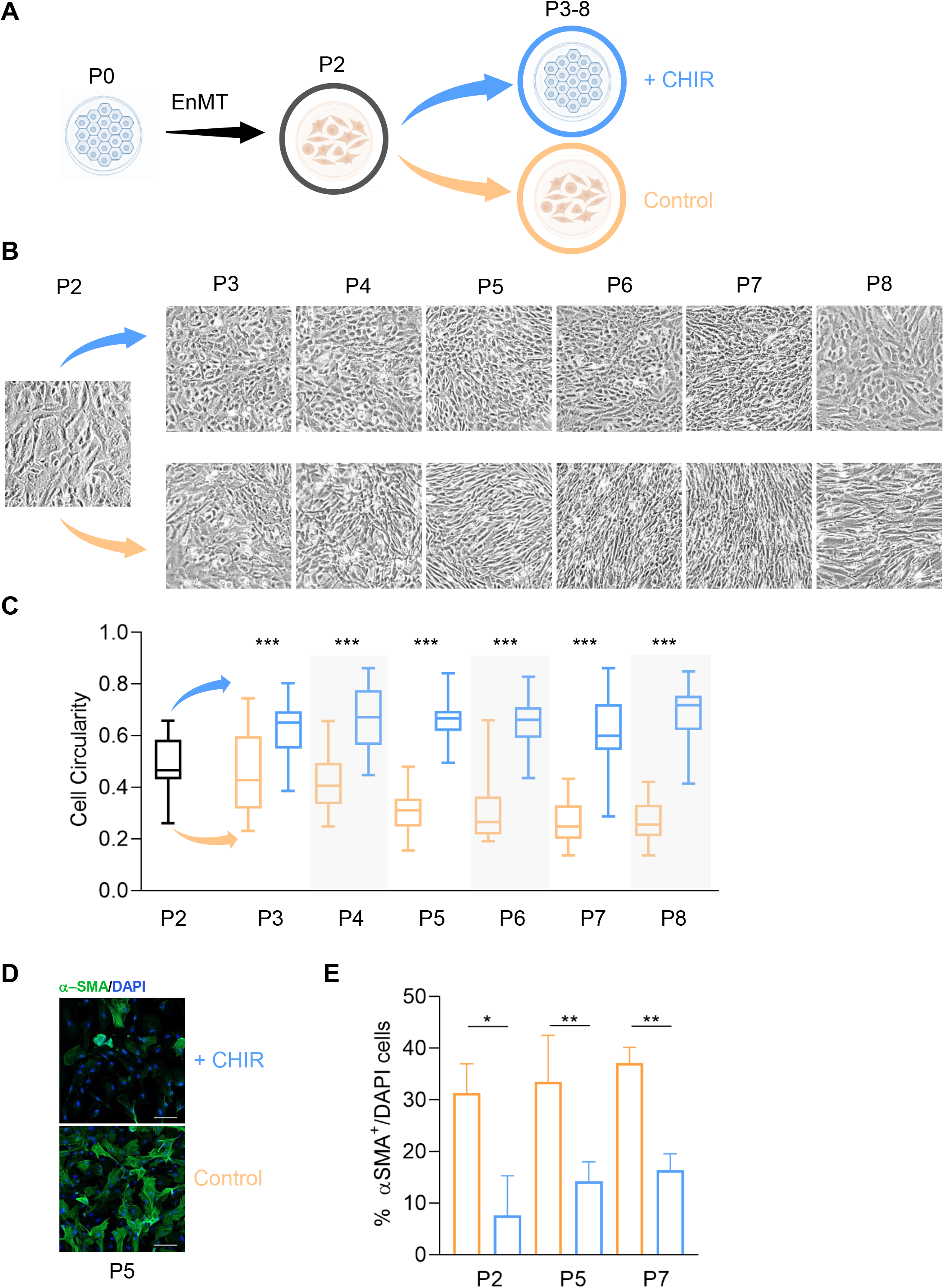
CHIR reverses the mesenchymal phenotype in primary cultures of HCEnCs. A) Schematic representation of the experimental plan for CHIR addition to HCEnCs culture, after EnMT. Image created with Biorender.com B) Representative phase-contrast images of primary HCEnCs treated with 1 μM CHIR at progressive passages (P3 to P8) following EnMT, showing the morphological differences with the untreated HCEnCs. C) Box plot comparing cell circularity values between HCEnCs treated with CHIR and their relative untreated control (n=3) at subsequent passages in culture. Cell circularity is expressed as values ± SD. D) Representative immunofluorescence microscopy images of HCEnCs at P5, showing expression of the α-SMA (green) EnMT marker on CHIR treated HCEnCs and their relative untreated control. DAPI (blue) counterstains nuclei. Scale bar 50 μm. E) Quantification of the percentage of HCEnCs expressing α-SMA protein (± SD) in samples treated with CHIR, compared with their relative untreated control, as seen in Figure 2D. Statistical significance was assessed using a Student t-test and set at p<0.05. p values are indicated as following: ns (not significative) when p>0.05, * when p<0.05, ** when p<0.01, *** when p<0.001.

A schematic diagram illustrates the experimental design for treating the spindle-shaped HCEnCs with CHIR from P3 to P8 (Figure 2A).

The morphological reversion of HCEnCs mesenchymal phenotype was observed in CHIR treated cells as compared to the control, at each passage in culture until P8 (Figure 2B). Cell circularity quantified the morphological transformation: a significant difference in cell circularity between CHIR and control samples was detected from the first CHIR application (P3, p=0.004) and maintained until P8 (p<0.0001) (Figure 2C).

CHIR treated samples presented a consistent decrease in α-SMA protein expression either at low or high passages (a representative image is shown for P5 in Figure 2D). Quantification of the immunofluorescence results showed that α-SMA expression was reduced in CHIR treated HCEnCs at P2, P5 and P7 to 7.6 ± 4.4%, 14.1 ± 1.9% and 16.3 ± 1.8% from its initial value of 31.3 ± 3.2%, 33.4 ± 4.5%, 37.2 ± 1.7%, respectively (Figure 2E).

The results obtained highlight that primary HCEnCs that underwent EnMT, if stimulated with CHIR, can reacquire their characteristic polygonal morphology and lose the α-SMA marker for several subsequent passages in culture.

### CHIR helps maintaining the correct expression and localization of HCEnCs markers

A deeper analysis of CHIR effects, protein expression and localization, as well as of multiple culture condition was conducted to try elucidating its mechanism on HCEnCs (Figure 3). A comparison between CHIR treated and untreated HCEnCs showed that several corneal endothelial markers such as Na^+^/K^+^ ATPase, ZO-1, β-catenin and N-cadherin were expressed correctly only in the CHIR treated HCEnCs, while they were absent and/or delocalised in the untreated control (Figure 3A). A caspase 3/7 assay was carried out in parallel to exclude the possibility of apoptosis events due to CHIR treatment. Figure 3B represents a comparison between CHIR previously treated and untreated HCEnCs the day after plating, before adding CHIR to the culture media (representative images obtained at P6). At any passage, HCEnCs appeared with a similar spindle shaped morphology at sub-confluency, which reverted to a polygonal morphology once at confluence if CHIR was added, while it remained elongated in the untreated control. To study if the CHIR effect could be exerted even at later passages, HCEnCs culture was further divided by adding two conditions to the CHIR-control at P6: in one CHIR was removed from the previously CHIR treated cells; while in the other CHIR was added to the cells that had not been treated in the previous passages (Figure 3C, P6 b). Whenever at confluence, the novel samples where CHIR was removed lost the polygonal morphology, while the control samples where CHIR was added only at P6 reacquired a polygonal morphology (a schematic is shown in Figure 3D). This tendency was confirmed by evaluation of cell circularity (Figure 3E): CHIR addition was able to revert the phenotype even at late passages, with HCEnCs presenting a circularity of 0.52, significantly different from the circularity of HCEnCs deprived of CHIR, equal to 0.26 (p<0.0001).

**Figure 3.**
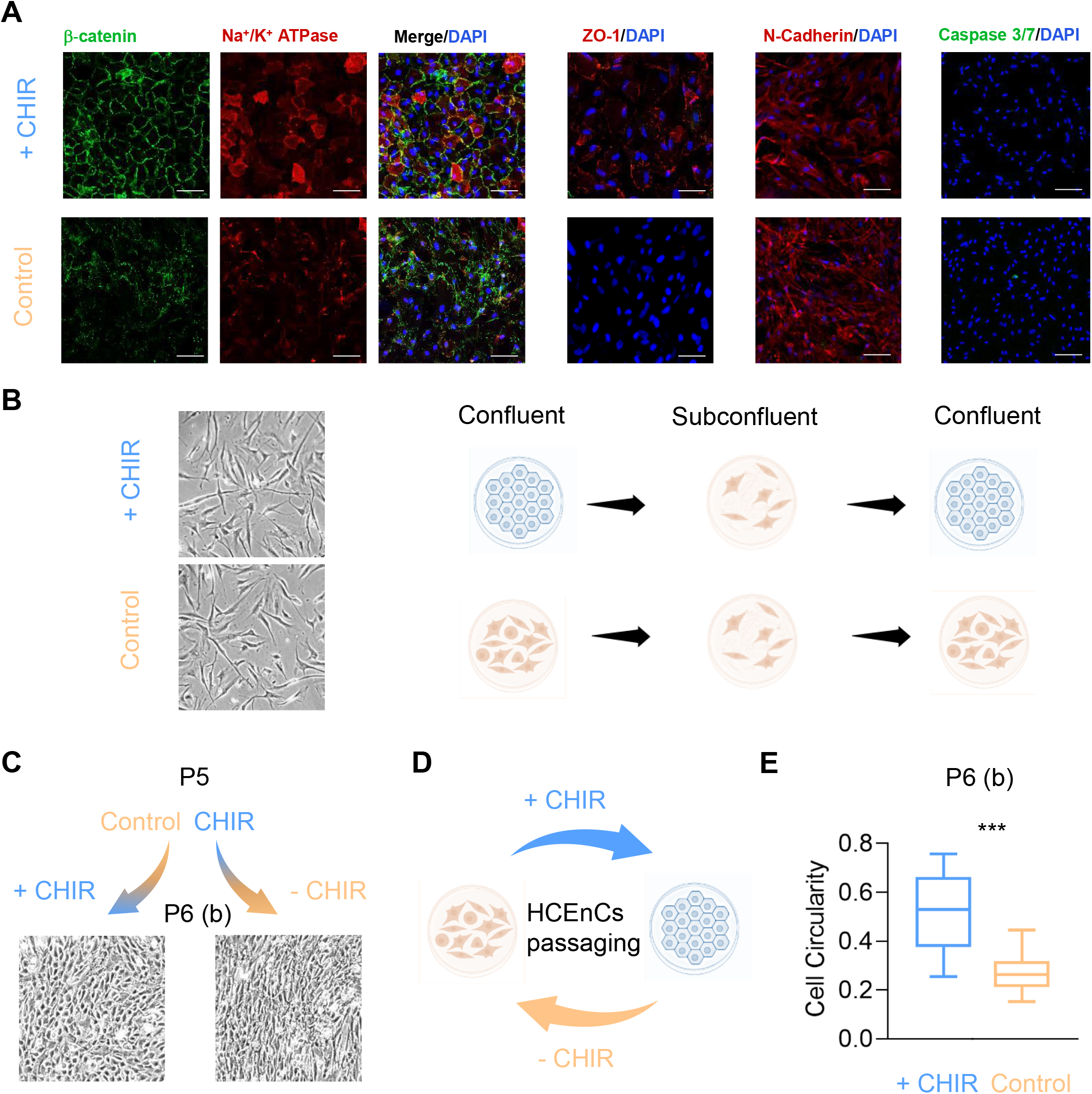
CHIR restores corneal endothelial protein expression and morphology at later stages. A) Representative immunofluorescence microscopy images showing expression of typical HCEnCs markers such as β-catenin (green, P7), Na^+^/K^+^ ATPase (red, P7), ZO-1 (red, P5), N-cadherin (red, P4) and Caspase 3/7 (green, P5) on CHIR treated HCEnCs and their relative untreated control. DAPI (blue) counterstains nuclei. Scale bar 50 μm, for Caspase 3/7 only scale bar 100 μm. B) Representative phase-contrast images of sub-confluent HCEnCs the day after plating, before being treated with CHIR (P6), showing no morphological differences with the untreated HCEnCs. A simplified scheme is shown on the right, done in Biorender.com. C) Representative phase-contrast images of HCEnCs untreated at P5 and treated with CHIR at P6, in parallel to HCEnCs treated with CHIR at P5, becoming untreated at P6. CHIR demonstrates to exert its effect also at a later stage, which is reversible upon passages. D) Schematic representation of the HCEnCs morphological change upon passaging when treated with CHIR, done in Biorender.com. E) Box plot comparing cell circularity values between HCEnCs treated with CHIR and their relative untreated control from Figure 3C. Cell circularity is expressed as values ± SD. Statistical significance was assessed using a Student t-test and set at p<0.05. p values are indicated as following: ns (not significative) when p>0.05, * when p<0.05, ** when p<0.01, *** when p<0.001.

Taken together, these results indicate how CHIR addition promotes the correct expression and localization of corneal endothelial characterizing proteins. At any passage, either in CHIR or control samples, cells cross an intermediate phase at subconfluence when they show an elongated morphology. The restoration of polygonal morphology can be induced by adding CHIR even at late passages in culture while it is abolished when, upon cell passaging, CHIR is removed.

### CHIR allows maintaining a polygonal phenotype in primary cultures of HCEnCs

Primary HCEnCs that still retained a polygonal morphology (P2), were treated with CHIR for several subsequent passages to evaluate the CHIR capacity to maintain the correct morphology and prevent the EnMT process (Figure 4).

**Figure 4.**
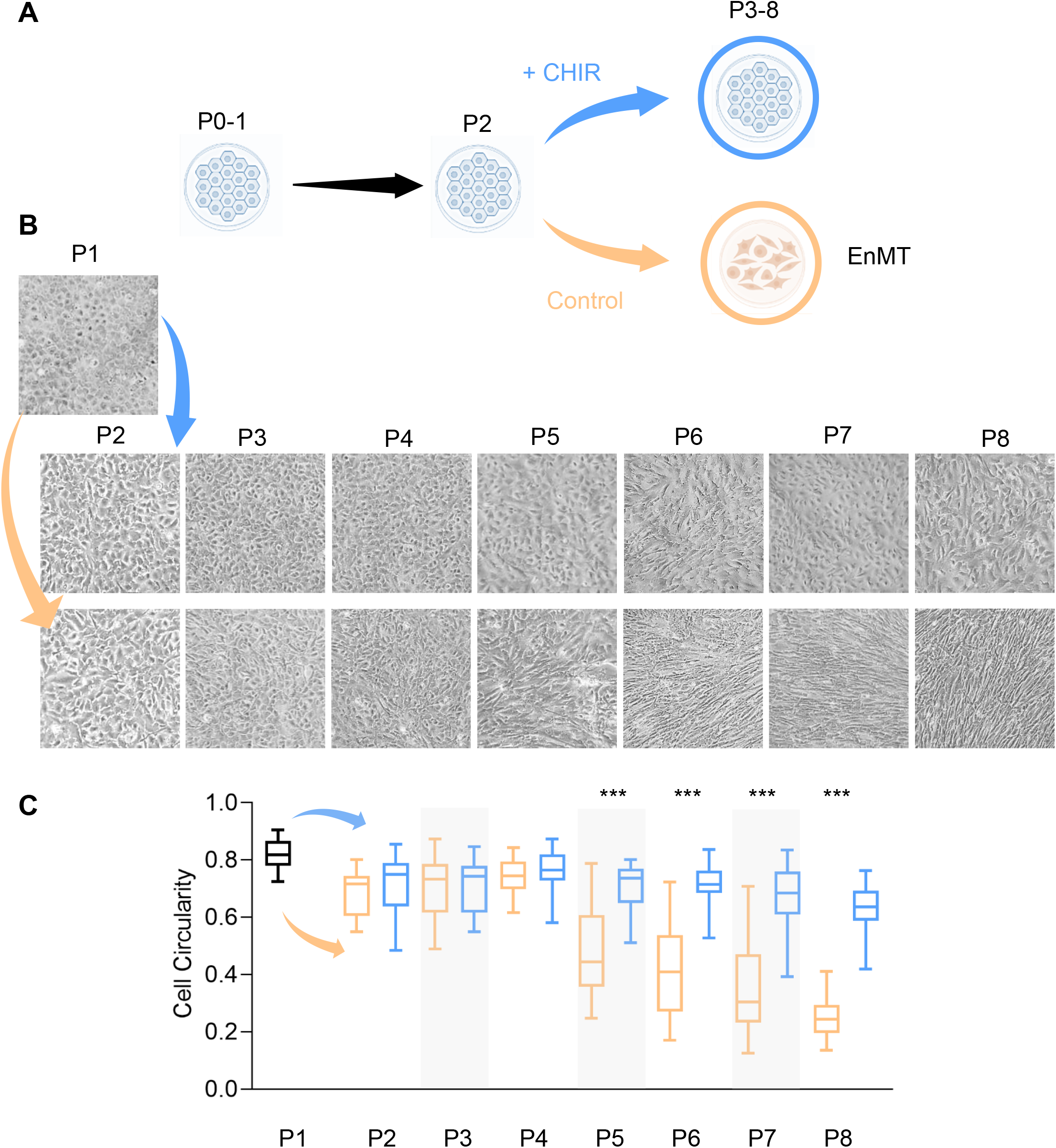
CHIR preserves the polygonal phenotype in primary cultures of HCEnCs. A) Schematic representation of the experimental plan for CHIR addition to HCEnCs culture, before EnMT. Image created with Biorender.com B) Representative phase-contrast images of primary HCEnCs culture at progressive passages (P3 to P8) treated with CHIR before EnMT, showing the morphological differences with the untreated HCEnCs. C) Box plot comparing cell circularity values between HCEnCs treated with CHIR and their relative untreated control (n=3) at subsequent passages in culture. Cell circularity is expressed as values ± SD. Statistical significance was assessed using a Student t-test and set at p<0.05. p values are indicated as following: ns (not significative) when p>0.05, * when p<0.05, ** when p<0.01, *** when p<0.001.

A scheme of the experimental design is shown in Figure 4A. CHIR was added to the culture media at any passage and compared with its relative untreated HCEnCs control: morphological differences started becoming evident at P3 and the discrepancy between the two conditions raised up as the passages increased (Figure 4B). Quantification of cell circularity revealed a significant difference between CHIR treated and untreated control (Figure 4C), starting from P5 (p<0.0001) and increasing progressively, until P8 (p<0.0001).

These data reveal how CHIR can prevent the loss of polygonal phenotype in primary HCEnCs for multiple passages in culture. This mechanism could be exploited to maintain HCEnCs in culture for a longer period of time, while they preserve the correct morphology.

### CHIR reduces cell proliferation rate in primary cultures of HCEnCs

Cell proliferation upon CHIR treatment was studied with multiple techniques (Figure 5). For the sample groups treated before or after acquiring an elongated morphology, the cells were counted upon confluency at any passage and compared between the CHIR treated and control HCEnCs (Figure 5A). The number of CHIR treated HCEnCs was constantly lower with respect to their relative untreated control, similarly in both sample groups. The following experiments on cell proliferation were carried out on the group sample that started treatment before changing morphology.

**Figure 5.**
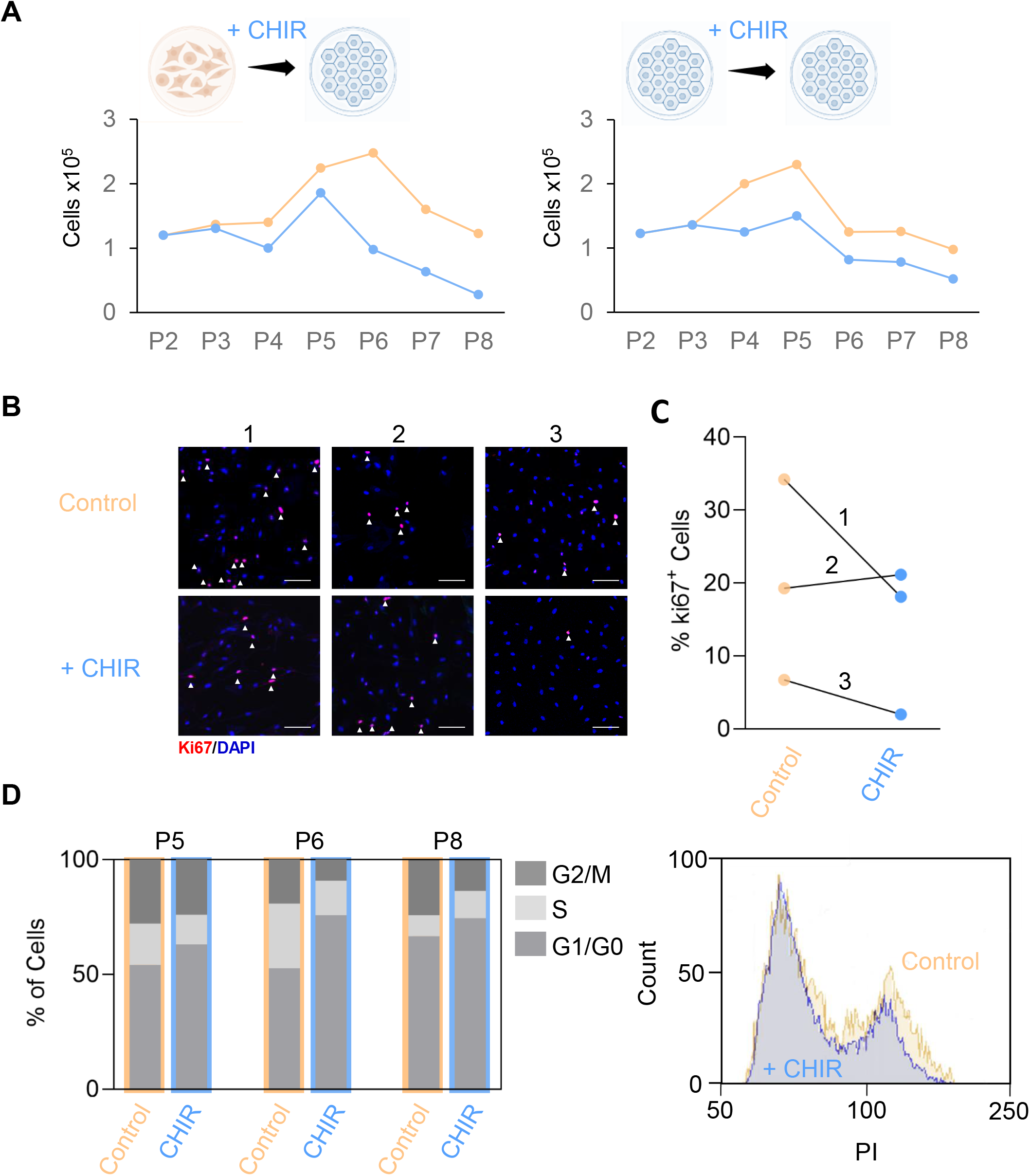
CHIR reduces cell proliferation rate in primary cultures of HCEnCs. A) HCEnCs count at progressive passages, compared between CHIR treated and untreated HCEnCs. Graphs show representative values for HCEnCs where CHIR treatment started before and after EnMT. B) Immunofluorescence microscopy images showing expression of ki67 (red) marker in sub-confluent HCEnCs at P5, untreated or treated with CHIR. DAPI (blue) counterstains nuclei. Scale bar 100 μm. C) Quantification of the percentage of HCEnCs expressing ki67 protein as seen in Figure 5B. D) Fluorescence Activated Cells Sorting (FACS) analysis of primary HCEnCs at P5-P6-P8 treated with CHIR and untreated control.

Immunofluorescence analysis on HCEnCs at P5 revealed an equal or lower amount of cells expressing the ki67 proliferation marker in CHIR treated cells, as evaluated in three different strains (Figure 5B and C).

A reduction in cell proliferation rate is confirmed by FACS analysis on sub-confluent HCEnCs cultures at high passages (P5 - P7 - P8), whenever the two conditions (CHIR-control) demonstrated a significant morphological difference between them at the previous passage upon confluency. In all the three passages analysed, CHIR enhanced the percentage of cells in G0/G1 phase of the cell cycle and concomitantly reduced the percentage of cells at G2/M (Figure 5D).

The data obtained demonstrate that CHIR addition in primary HCEnCs cultures inhibits the cell proliferation rate, as evaluated with different assays.

### CHIR effects on gene expression in primary cultures of HCEnCs

A set of selected genes were analysed at RNA level to get novel insights into the mechanism of CHIR effect on preserving HCEnCs morphology and protein expression (Figure 6). At low passages, none of the selected genes showed any difference between the CHIR treated and untreated HCEnCs but we could appreciate a significant up-or down-regulation at high passages (P6-P7) for the following genes.

**Figure 6.**
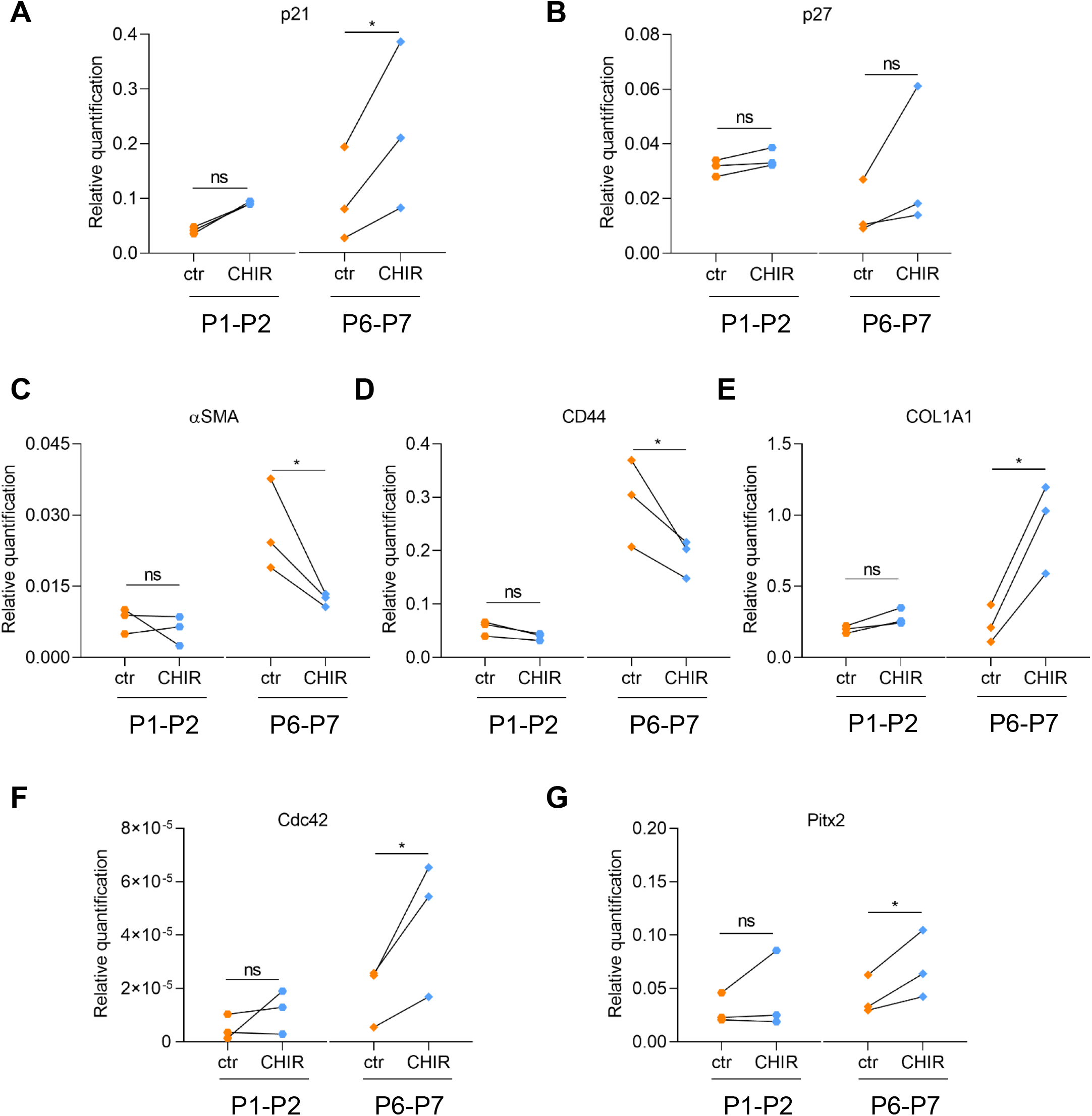
CHIR effects on gene expression in primary cultures of HCEnCs. Panel of genes (p21, p27, α-SMA, CD44, COL1A1, CdC42, Pitx2) whose mRNA expression has been evaluated on primary HCEnCs at low (P1-P2) and high (P6-P7) passages by quantitative (q) RT-PCR. mRNA levels for each gene are shown as relative expression ± SD. Statistical significance was assessed using a Student t-test (Ratio paired) and set at p<0.05. p values are indicated as following: ns (not significative) when p>0.05, * when p<0.05, ** when p<0.01, *** when p<0.001.

At high passages, genes related to HCEnCs cell cycle such as p21 and p27 have been investigated: CHIR stimulation induced an increased expression of p27 (p=0.06) and p21 (p=0.02), the latter resulting significant if compared with untreated cells (Figure 6A and 6B). EnMT marker genes were found here significantly downregulated at RNA level upon CHIR treatment: α-SMA (p=0.03) and CD44 (p=0.04). This data is in accordance with the CHIR induced EnMT reduction, supported by a concomitant maintenance of hexagonal cell morphology and a decreased α-SMA protein expression, as shown above.

Surprisingly COL1A1, an important mediator of tissue wound healing^42^, resulted upregulated at high passages following CHIR induced β-catenin stabilization (p=0.01). Cdc42, a marker of the non-canonical Wnt pathway regulating HCEnCs migration^34^, and Pitx2, a protein involved in the development of the cornea^43^, were significantly upregulated in CHIR treated cells (p=0.01 and p=0.03, respectively).

Additional genes such as SOX2, involved in wound healing in CE^44^, and Zeb1, found to mediate fibrosis in CE^45^, were assessed herein and compared between CHIR treated and untreated cells but showed no significant differences and low levels of expression either at low or high passages (data not shown).

Expression analysis of selected genes confirmed an increased expression of cell cycle inhibitors, that well matches the reduced proliferation (Figure 5) as well as the reduced EnMT associated markers. Evaluation of other genes revealed possible new perspectives in the pathways that regulates HCEnCs growth and barrier formation.

### β-catenin signalling intersects with TGFβ pathway on HCEnCs

Further analysis has been carried out on HCEnCs to better delineate the intracellular pathway regulation of β-catenin and TGFβ signalling upon CHIR and TGFβ Inhibitor (TGFβI) treatments (Figure 7).

**Figure 7.**
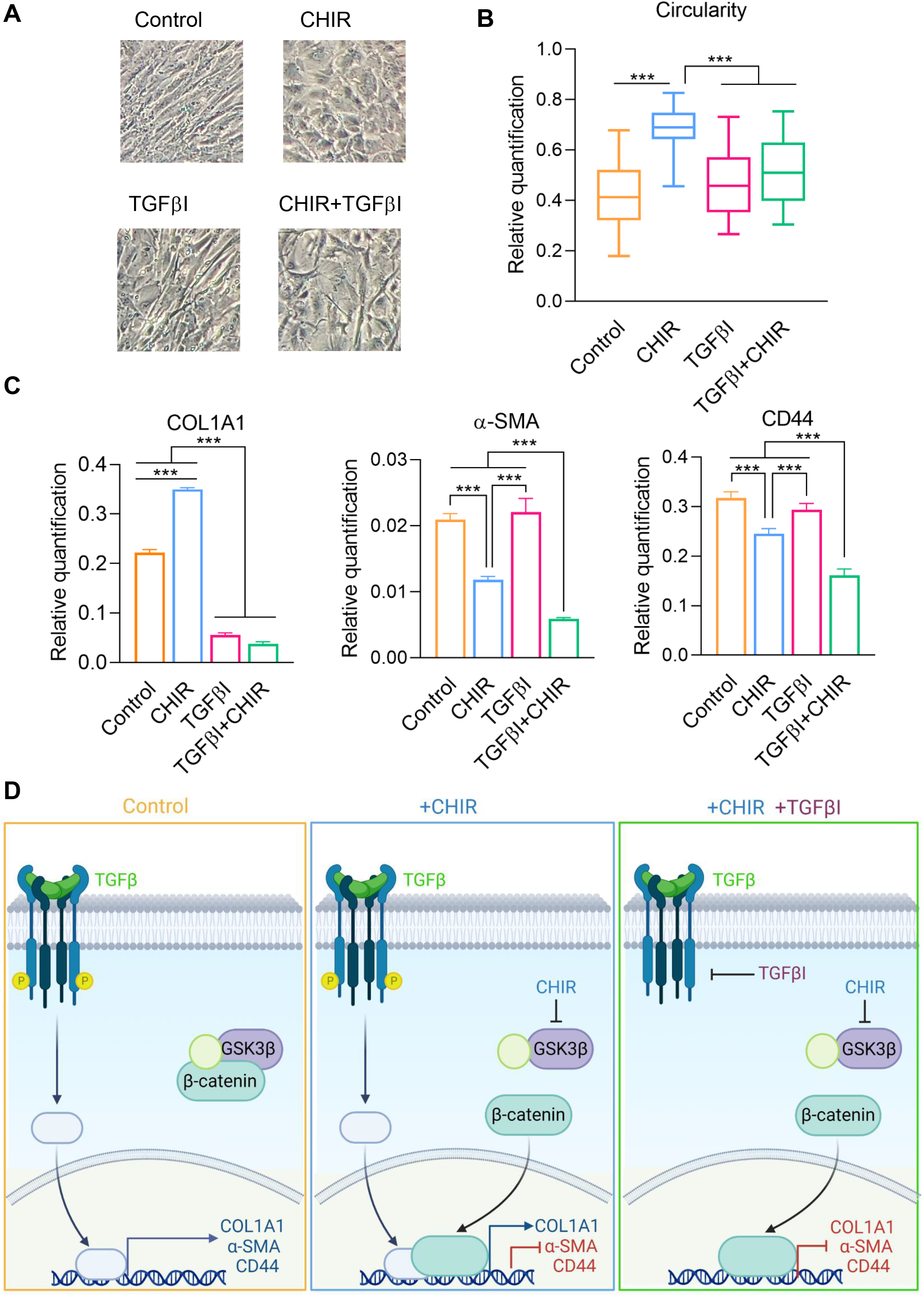
β-catenin signalling intersects with TGFβ pathway on HCEnCs. A) Representative phase-contrast images of primary HCEnCs culture at P4 treated with CHIR, TGFβI and CHIR+TGFβI showing the morphological differences with the untreated HCEnCs. B) Box plot comparing cell circularity values between HCEnCs treated and untreated cells at subsequent passages in culture. Cell circularity is expressed as values ± SD. C) mRNA expression of COL1A1, α-SMA and CD44 of primary HCEnCs culture at P4 (± SD). Statistical significance was assessed using a Student t-test (Ratio paired for the RT-PCR analysis) and set at p<0.05. p values are indicated as following: ns (not significative) when p>0.05, * when p<0.05, ** when p<0.01, *** when p<0.001. D) Schematic representation of a putative molecular pathway based on our finding, involving β-catenin and TGFβ. Image created with Biorender.com

A morphological variation was assessed in HCEnCs at different passages: it was evident how CHIR preserved the hexagonal morphology, while the TGFβI and CHIR+TGFβI conditions did not show any improvement from the untreated control (Figure 7A shows representative images at P4). Quantification of cell circularity at P4 revealed a significant difference between CHIR treated and TGFβI or CHIR+TGFβI cells (p<0.0001); while the latter two conditions did not appear different from the control (Figure 7B).

Gene expression revealed that TGFβI treatment significantly decreases COL1A1 expression in HCEnCs (p<0.0001), as known from literature and similarly to what observed with the combined effect of CHIR+TGFβI (p<0.0001). While inhibition of the TGFβ signalling did not result in a significant variation of α-SMA and CD44 expression, the cumulative effect of CHIR and TGFβI determined a significant downregulation of the two EnMT markers (p<0.0001).

The results obtained herein suggest that TGFβ and β-catenin act in a coordinated fashion to regulate HCEnCs expansion.

## Discussion

The visual stimuli, fundamental for human quality of life, pass through a clear cornea. While the outer corneal epithelium is constantly renewed by stem cells^46^ and advanced cell therapy are already a clinical practice^47^, regenerative medicine approaches are still at their debut for the inner CE^8^.

HCEnCs, although arrested in G1 phase of the cell cycle *in vivo*^*48*^, retain a limited proliferative capacity *in vitro* that allows a contained expansion in culture. This is only possible for a short number of passages since HCEnCs easily undergo EnMT and lose the characteristic morphology and expression markers, required for their function^9^.

During EnMT, HCEnCs acquire an elongated shape and EnMT markers such as α-SMA^13^, while losing cell-cell contact and typical corneal endothelial markers such as ZO-1 and Na^+^/K^+^ ATPase^49^. Several attempts have been made in preventing EnMT in HCEnCs culture through growth medium optimization^22, 28^ but no reports to date have thoroughly described a method for reverting the mesenchymal phenotype of HCEnCs for subsequent passages in culture, once they have undergone EnMT.

CHIR99021, a GSK-3β inhibitor that induces β-catenin activation, demonstrated capable of reverting the elongated shape in HCEnCs primary culture by reducing α-SMA expression and EnMT, while promoting the expression of characteristic CE markers and the formation of cell-cell junction upon confluency. Regulation of β-catenin was previously found to have a pivotal role in CEnCs expansion, since β-catenin related effectors were over-expressed in cells *in vitro* when compared with the corresponding corneal explants^16^. This was partially explained by experiments proving how cellular proliferation of CEnCs was impaired by blocking β-catenin pathway (using quercetin). Although CHIR stimulation promoted β-catenin stability, it did not induce further proliferation^16^. Nevertheless, the involvement of β-catenin pathway in other mechanisms of HCEnCs during cell expansion, such as EnMT^13^, has been proposed^13, 16^, although it remains largely unexplored. The intracellular function of β-catenin is tissue and context specific and its interaction with different nuclear transcription factors may promote alternative cellular processes^29, 50^. Of particular interest, accumulating evidence indicates how this effector may push not only towards proliferation^51-52^ but also toward differentiation^53-55^. Moreover, β-catenin is also a key regulator of mesenchymal transformation during development^56^, fibrosis^57-58^ and wound healing^59^. In accordance with this, we have demonstrated here that, when over-stimulated with 10 μM CHIR, β-catenin promotes EnMT; otherwise, when more finely tuned (1 μM CHIR), it blocks and reverts EnMT in primary HCEnCs for several subsequent passages in culture, also in old donors. In particular, once CHIR was added to HCEnCs that had already underwent EnMT, cell morphology, as measured by cell circularity^22^, was maintained through the passages up to passage 8, while α-SMA was significantly downregulated both at RNA and protein level and CD44, another marker of EnMT^60^, was significantly downregulated at RNA level. CEnCs markers such as Na^+^/K^+^ ATPase, N-cadherin and ZO-1^61^, were also preserved upon CHIR treatment. The same result was obtained by adding CHIR in HCEnCs that still maintained a normal hexagonal phenotype. In accordance with the observed ability of promoting barrier formation while reverting the EnMT process, β-catenin retains a role in stabilizing the tight junctions by increasing ZO-1 expression^62^.

All together, these results suggest how EnMT may be reversed, at least until a certain limit. Consistently with this observation, sub-confluent HCEnCs culture presented a slight elongated phenotype, both in control and CHIR treated cells, independently from the phenotype they reacquired when confluent, either hexagonal or elongated. These data suggest that HCEnCs may undergo a transient mesenchymal transformation when proliferating and a subsequent mesenchymal-to-endothelial (MEnT) transformation when reaching confluency. Although some observations reported here are only hints of this process, this aspect should be further confirmed and investigated to identify possible key regulators orchestrating this reversible mechanism.

As we previously noticed^16^, treatment with CHIR did not stimulate proliferation nor induced cell death; on the contrary it slightly decreased cells entering G2/M and consequently actively proliferating, probably through the activation of cell cycle inhibitors p21 and p27 that we found significantly overexpressed in CHIR treated cells. This result is in contrast with what reported by Wang *et al*. in a corneal endothelial cell line (B4G12), where CHIR stimulation promoted an increased proliferation^39^. However, the B4G12 cell line is not comparable to a human corneal endothelial primary culture as presenting a different pattern of marker expression, and an altered cellular proliferation mechanism is typical in immortalised cell lines^63-64^.

Furthermore, here we found how the intersection of β-catenin pathway with that of TGF-β regulates Collagen I expression. Although also described as a marker of EnMT^14^, Collagen I is involved in tissue wound healing, a process that requires extracellular matrix deposition and remodelling^42^. TGF-β is a well known regulator of Collagen I expression^65^ and here we confirmed that, by inhibiting this pathway, Collagen I expression was severely down-regulated. Conversely, CHIR stimulation promoted an increase of Collagen I expression, possibly because the interaction of both (TGFβ and β-catenin) pathways is necessary to stimulate Collagen I production. On the other hand, inhibition of TGFβ did not decrease α-SMA and CD44 expression, nor reverted cellular morphology toward a hexagonal shape, while CHIR stimulation was able to promote the latter processes associated with EnMT. A combination of CHIR and TGFβI induced in part a cumulative effect: it reduced Collagen I expression and further decreased expression of α-SMA and CD44. However, there was not any improvement on the cellular shape if compared with the CHIR condition, suggesting that the intersection of TGFβ and β-catenin is important in HCEnCs for maintenance of the correct morphology.

The cumulative downregulation of α-SMA and CD44 might be caused by a sustained regulatory effect of TGFβ downstream effectors (Smad) on β-catenin function. Intriguingly, this combined effect is in contrast with reports on other cellular types^66^, confirming the context specificity of the β-catenin regulated processes. Altogether, these results support the hypothesis of a crosstalk between TGFβ and β-catenin pathways that is responsible for CEnCs fate, although the combined effect caused by a concomitant dysregulation should be further dissected, considering different dosages and timings. To define whether up-regulation of the β-catenin pathway may have an impact on indirect pathways also we analysed the expression of other effectors, selected based on previous findings in CE. Although not highly expressed in HCEnCs, Cdc42, a downstream target of the non-canonical Wnt pathway resulted significantly upregulated at high passages following CHIR treatment, as previously observed in B4G12 cell line^39^. However, while in B4G12 it was associated to an increased proliferation, herein it correlates with a decreased cell growth, similarly to what previously found in neural stem cells^67^.

Pitx2, a homeobox gene involved in corneal development through Wnt interaction^43^, was found here upregulated upon β-catenin stabilization, as previously observed in HCEnCs by Hatou *et al*.^38^, revealing a possible role for this gene in supporting β-catenin for determining cell fate.

Collectively, this work identified CHIR99021 as a single factor able to revert the elongeted phenotype of HCEnCs for several subsequent passages in culture. CHIR99021, through specific GSK-3β inhibition and the consequent cytoplasmic β-catenin stabilization, promotes corneal endothelial differentiation from an elongated to a hexagonal phenotype. HCEnCs pass through a transient EnMT during expansion in culture (sub-confluence) that is reverted at confluency only if β-catenin is properly dosed, in order to induce the correct phenotype and junction formation. Thus, based on the findings of this study, CHIR99021 inhibits EnMT in corneal endothelium through β-catenin stabilization, which interacts with other pathways such as TGFβ for determining cell differentiation.

## CONCLUSION

The identification of CHIR as a single factor able to revert the HCEnCs elongated phenotype in culture represents a substantial achievement for maintaining HCEnCs morphology and markers *in vitro* until late passages.

Further investigations are necessary for unravelling the precise mechanisms involved in HCEnCs proliferation and differentiation, and how Wnt/β-catenin pathway interplays with others, such as TGFβ’s. These findings would lay the path for developing therapeutic strategies *in vivo* using compounds that maintain corneal endothelial differentiation, such as CHIR, alone or in combination with other factors that promote a controlled cell proliferation.

## Acknowledgments

The authors would like to thank the students that helped within this project.

